# Functional Evidence for Biased Inhibition of G protein Signaling by YM-254890 in Human Coronary Artery Endothelial Cells

**DOI:** 10.1101/2020.08.12.248468

**Authors:** Qianman Peng, Saud Alqahtani, Mohammed Zahid A Nasrullah, Jianzhong Shen

**Author notes:** To whom correspondence should be addressed: Jianzhong Shen, Ph.D., Associate Professor, Department of Drug Discovery and Development, Harrison School of Pharmacy, Auburn University, Pharmaceutic Research Building, 720 S. Donahue Dr., Auburn, AL 36849.

## Abstract

Small molecular chemicals targeting individual subtype of G proteins including Gs, Gi/o and Gq has been lacking, except for *pertussis toxin* being an established selective peptide inhibitor of the Gi/o protein. Recently, a cyclic depsipeptide compound YM-254890 isolated from culture broth of *Chromobacterium sp*. was reported as a selective inhibitor for the Gq protein by blocking GDP exchange of GTP on the α subunit of Gq complex. However, functional selectivity of YM-254890 towards various G proteins was not fully characterized, primarily due to its restricted availability before 2017. Here, using human coronary artery endothelial cells as a model, we performed a systemic pharmacological evaluation on the functional selectivity of YM-254890 on multiple G protein-mediated receptor signaling. First, we confirmed that YM-254890, at 30 nM, abolished UTP-activated P2Y2 receptor-mediated Ca^2+^ signaling and ERK1/2 phosphorylation, indicating its potent inhibition on the Gq protein. However, we unexpectedly found that YM-254890 also significantly suppressed cAMP elevation and ERK1/2 phosphorylation induced by multiple Gs-coupled receptors including β_2_-adrenegic, adenosine A_2_ and PGI_2_ receptors. Surprisingly, although YM-254890 had no impact on CXCR4/Gi/o protein-mediated suppression of cAMP production, it abolished ERK1/2 activation. Further, no cellular toxicity was observed for YM-254890, and it neither affected A23187- or thapsigargin-induced Ca^2+^ signaling, nor forskolin-induced cAMP elevation and growth factor-induced MAPK signaling. We conclude that YM-254890 is not a selective inhibitor for Gq protein; instead, it acts as a broad spectrum inhibitor for Gq and Gs proteins and exhibits a biased inhibition on Gi/o signaling, without affecting non-GPCR-mediated cellular signaling.

---

G protein-coupled receptor (GPCR) has been the primary therapeutic targets for many diseases. Agonists and antagonists of various GPCR are estimated to occupy approximately 35% of the drug market (1). Extracellular signals perceived by GPCR are transmitted via G proteins and trigger intracellular signaling cascades resulting in a plethora of physiological responses. G proteins usually are categorized into four main classical subfamilies according to their α subunits: Gα_q/11_, Gα_s_, Gα_i/o_, Gα12/13 (2). Main subunits Gα_q/11_, Gα_s_, Gα_i/o_ and G_12/13_ are commonly thought to be coupled with phospholipase C (PLC), adenylyl cyclase activation, adenylyl cyclase inactivation, and other small GTPase families, respectively (3). Gα_q/11_ activates phospholipase Cβ pathway leading to intracellular Ca^2+^ mobilization. Gα_s_ and Gα_i/o_ proteins regulate adenylyl cyclase activation and inhibition, respectively, which control intracellular cAMP level. Thus, G proteins are important signal transducing molecules for various cellular responses. Malfunction of GPCR signaling pathways are involved in many diseases, such as diabetes, cardiovascular diseases, and certain forms of cancers (4).

Even though there are roughly 800 human genes coding for various GPCRs and about 300 of them have been identified as real or potential pharmaceutical targets (5), the modern techniques to analyze the intracellular signaling network for GPCR is still limited. Using G protein inhibitor is probably one of the most straightforward ways to analyze the intracellular signaling pathways induced by GPCR of interest. However, only two types of specific G protein inhibitors were identified and developed at present. *Pertussis toxin* (PTX) is the only specific Gi/o inhibitor widely used in the research community for decades with virtually no controversy (6). However, similar inhibitors specifically targeting other types of G proteins have been very limited. Recently a cyclic depsipeptide compound YM-254890, which was originally isolated from *Chromobaterium* sp. and claimed as the first specific Gα_q_ subfamily inhibitor (7, 8), was released and available on the market. After YM-254890, another G protein inhibitor UBO-QIC, also termed as FR90035, was developed based on the structure of the YM compound (9, 10). Except for these, there are no other types of specific G protein inhibitors available on the market.

YM-254890 has been shown to specifically inhibit cell signaling mediated by the Gα_q_-coupled receptors with a high potency (IC50 around 30 nM). It was reported that YM-254890 inhibits Gα_q_-coupled GPCR signaling by blocking GTP exchange from GDP, thus preventing Gα_q/11_ activation (11). Also, it was shown to inhibit calcium mobilization induced by various Gα_q_-coupled receptors including the P2Y1, P2Y2 purinergic receptors and the CysLT-R1/R2 receptors, without affecting P2X and L-type Ca^2+^ channel-induced Ca^2+^ influx (8). Furthermore, studies indicated that YM-254890 did not affect cAMP level that is decreased by activation of the G_i_-coupled P2Y12 receptor (8). Altogether, these findings suggested that YM-254890 might function as a selective and specific inhibitor for the G_q_ protein in the past decade.

However, it should be noted that studies of YM-254890 have been limited to few GPCRs and focused on limited number of cell signaling readouts such as intracellular Ca^2+^ mobilization and change of cAMP levels. Given that many receptor signaling pathways including the MAPK pathways are not necessarily coupled with the Ca2^+^ and cAMP second messenger systems, and that some GPCR can initiate signal transduction that are independent of G proteins (12), we thought it is important to study the pharmacological effects of YM-254890 on receptor-induced signaling pathways beyond Ca^2+^ and cAMP. Here, we provided strong evidence that YM-254890 is not a selective inhibitor for the Gq proteins; instead, it acts as a broader spectrum inhibitor for G_q_ and G_s_ with a unique biased inhibition on the Gi/o proteins, without affecting non-GPCR-mediated cellular signaling.

## EXPERIMENTAL PROCEDURES

### Cell culture

HCAEC were cultured in EBM-2 medium supplemented with VEGF, FGF, EGF, IGF, ascorbic acid, GA1000 (Lonza), and 5% FBS at 37°C in a humidified atmosphere of 5% CO_2_ as we previously reported (13, 14). HCAEC were used between the third and eighth passages. Before stimulation, cells were seeded to grow for 24h and starved overnight. Where inhibitor or antagonist was used, cells were pretreated with the inhibitor/antagonist for 40 minutes before cell stimulation.

### Intracellular Ca^2+^ mobilization assay

Cells were seeded at a density of 4×10^4^ cells per well into 96-well culture plates and cultured for one day. On day two, the original medium was removed and the assay medium from FluoForteTM kit (Enzo Life Sciences) containing the Ca^2+^ dye was added and receptor-mediated Ca^2+^ mobilization was determined as previously described (13). Fluorescence was determined immediately after adding of different reagents, with an excitation wavelength set to 485 nm and an emission wavelength set to 525 nm, and readings were taken every 1s for 500s. For antagonist inhibition experiment, cells were pre-incubated with the antagonist for 45 min before agonist addition. Measurement of Ca^2+^ signal was performed with the fluorometer plate reader (BMG FLUOstar) with a 490/525nm band-pass filter, the results of which was shown as relative fluorescence units (RFU).

### Western blotting assay

After stimulation, cells were lysed, and standard Western blotting assay was performed as previously described (14, 15). The individual primary antibodies used were anti-p-ERK1/2, anti-p-p38, anti-p-SAPK/JNK, anti-ERK1/2, anti-p38, anti-SAPK/JNK (Cell Signaling Technology). Equal protein loading was verified by stripping off the original antibodies and re-probing the membranes with a different primary antibody such as anti-GAPDH (Cell Signaling Technology). *Pertussis toxin* (PTX) treatment (100 ng/ml, 4h) was used to suppress Gα_i/o_ activity, after which cells were stimulated and lysed for Western blotting assay.

### Intracellular cAMP measurements

Intra-cellular cAMP levels were determined by using the Direct cAMP Elisa kit (Enzo Life Sciences) as previously reported (16). HCAEC were grown to confluence in 12-well plates and starved for overnight. Cells were pretreated with YM-254890 (30 nM, 45 min) to inhibit G protein of interest, and Gi protein-mediated cAMP suppression was achieved by stimulation of the CXCR4 receptor by SDF-1(100ng/mL) in the presence of forskolin. During the treatment, 10 μM Rolipram was used to inhibit cellular phosphodiesterase activity. In another set of experiments, accumulation of cellular cAMP was measured after stimulation of the cells with 10 μM forskolin or 100 μM adenosine. After 5min, the incubation was terminated by pouring off the medium and adding 300 μL HCl (0.1M). After cell lysis in the 37°C incubator, the samples were centrifuged (1000g × 5min) and the supernatants were acetylated as instructed by the kit. 100 μL of the acetylated supernatant was used for the Elisa assay followed the protocol of the kit. cAMP levels were normalized for the protein content of each sample as determined by the Bradford assay.

### Cell toxicity assay

HCAEC were seeded at a density of 4×10^4^ cells per well into 96-well culture plates and cultured for one day. On day two, the original medium was replaced by fresh culture media. 10% volume of resazurin (R&D Systems) was added into 96-well culture plates, and YM-254890 at different concentrations were then added into culture media and resazurin mix system. After 1h drug treatment, fluorescence was detected immediately with an excitation wavelength set to 544 nm and an emission wavelength set to 590 nm as we previously reported (15). Measurement of the signal was performed with the fluorometer plate reader (SpectraMax M5e) with a 544 /590nm band-pass filter, the results of which were shown as relative fluorescence units (RFU). Hydrogen peroxide was used as a positive control.

### Materials

Dulbecco’s Modified Eagle medium (DMEM) and EBM-2 were purchased from Lonza. FluoForte™ Kit was purchased from Enzo Life Sciences. Fetal Bovine Serum (FBS) was purchased from Thermo Fisher Scientific. DNA primers were purchased from Integrated DNA Technologies. YM-254890 was purchased from Adipogen. Other reagents including UTP and ATP were obtained from Sigma.

### Data analysis

All data were analyzed by Prism 5 (Graphpad Software Inc.). Data are expressed as the mean ± S.E.M. The means of two groups were compared using Student’s t-test (unpaired, two-tailed), and one-way analysis of variance was used for comparison of more than 2 groups with p<0.05 considered to be statistically significant. Unless otherwise indicated, all experiments were repeated at least three times.

## RESULTS

### Effect of YM-254890 on G_q_-coupled P2Y2R-induced Ca^2+^ signaling

To confirm whether YM-254890 (Fig. 1A) is a Gαq protein inhibitor, we examined the effect of YM-254890 on Ca^2+^ mobilization induced by the Gq-coupled P2Y2 receptor (P2Y2R) in HCAEC. Our previous study confirmed that the P2Y2R is the only UTP/ATP-sensitive nucleotide receptor expressed in HCAEC (13). As shown in figure 1, stimulation of the endothelial P2Y2R by UTP or ATP all triggered a peak Ca^2+^ response as anticipated. Pretreatment of the cells with YM-254890 at 30 nM, a dose around its reported IC_50_ (~50 nM) (14), abolished the Ca^2+^ response induced by the maximal dose of UTP and ATP (100 μM) (Fig. 1B, 1C). These results confirmed prior studies indicating that YM-254890 functions as a potent Gα_q_ protein inhibitor.

**Figure 1.**
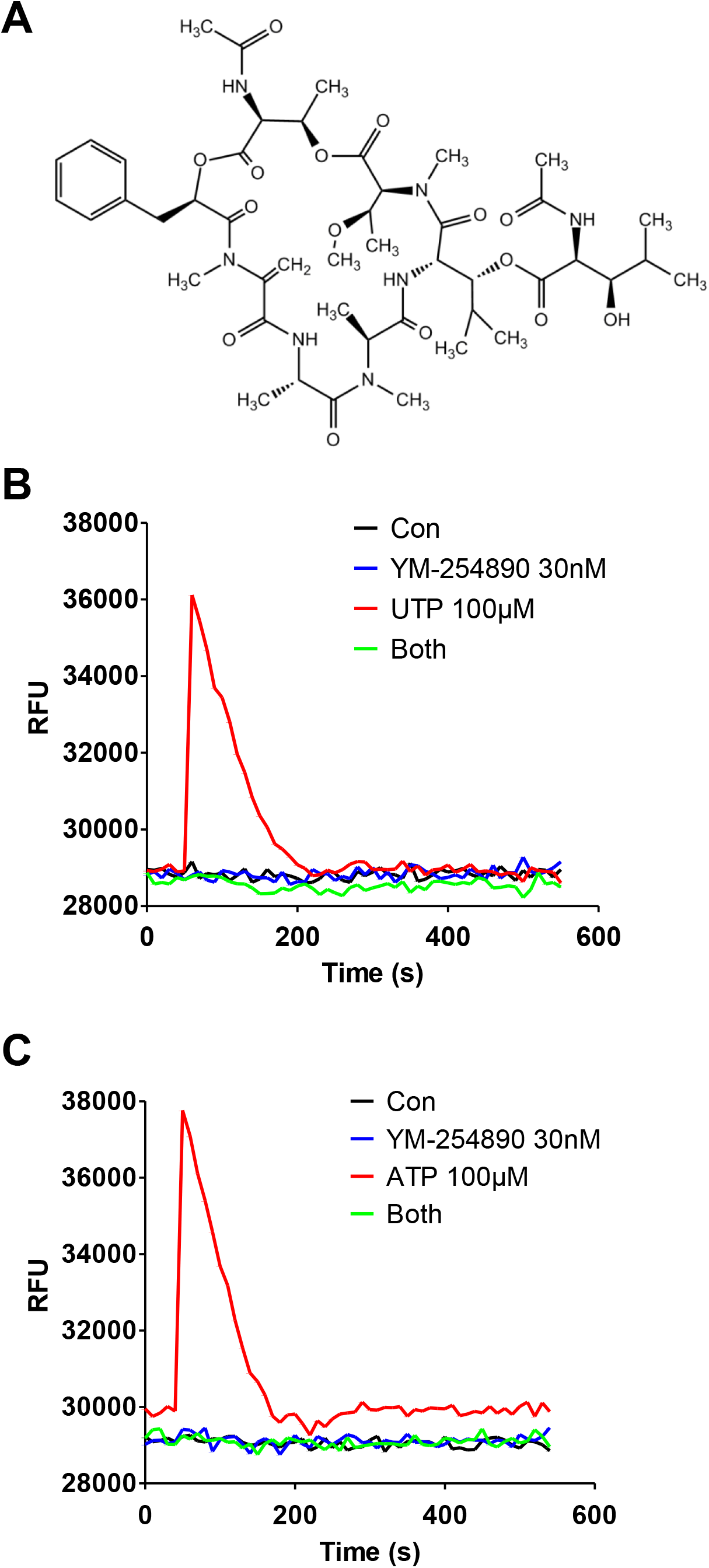
Effect of YM-254890 on G_q_-coupled P2Y2R-induced Ca^2+^ signaling in HCAEC. P2Y2R agonist UTP (100 μM)- or ATP (100 μM)-induced intracellular [Ca^2+^] mobilization was determined in HCAEC after pretreatment of the cells for 40 min with 30 nM YM-254890 (structure shown in **A**). YM-254890 abolished UTP-induced (**B**) and ATP-induced (**C**) [Ca^2+^] increase in HCAEC. Measurement of Ca^2+^ signal was performed by a fluorometer plate reader with a 490/525nm bandpass filter and Fura-4 as the probe, the results of which were shown as relative fluorescence units (RFU). Tracings shown are representative data of three independent experiments.

### Effect of YM-254890 on G_i/o_-coupled CXCR4 receptor-mediated cAMP inhibition

Since previous study reported that YM-254890 did not affect cAMP level that was inhibited by activation of the G_i/o_-coupled P2Y12 receptor expressed in C6-15 cells (8), we were curious to know if this selectivity could be achieved in other G_i/o_-coupled receptors. Here we chose the G_i/o_-coupled chemokine receptor CXCR4 as a testing target. Figure 2 shows that stimulation of the endothelial CXCR4 receptor by its natural ligand SDF-1 (CXCL12) caused ~40% inhibition of forskolin-induced cAMP elevation as expected. However, this significant inhibitory effect on cAMP production was not affected by 30 nM YM-254890 pre-treatment (Fig. 2), suggesting that consistent with prior study, YM-254890 is not a Gαi protein inhibitor.

**Figure 2.**
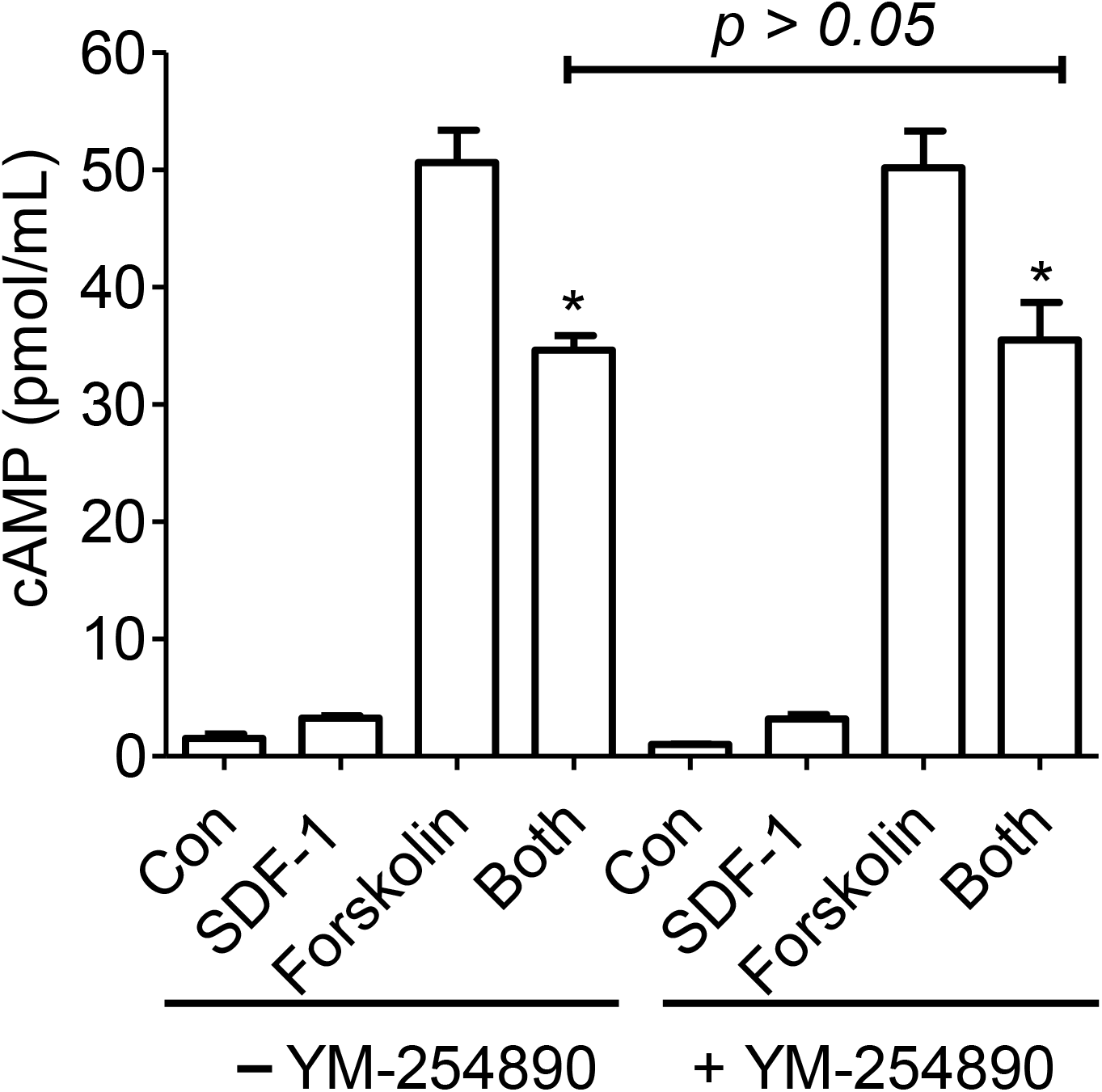
Effect of YM-254890 on G_i_-coupled CXCR4 receptor-mediated cAMP inhibition in HCAEC. Stimulation of G_i_-coupled CXCR4 receptors by SDF-1 (100 ng/mL) resulted in an inhibition of forskolin (10 μM)-induced cAMP elevation in HCAEC. This inhibitory effect on cAMP production was not affected by 30 nM YM-254890 pre-treatment for 40 min. cAMP assays were performed by ELISA as detailed in “Materials and methods” and data are the mean ± SEM from three independent experiments performed in triplicate. * *p*< 0.05 relative to the respective forskolin groups.

### Effect of YM-254890 on G_s_-coupled receptor-induced cAMP elevation

To further examine the specificity of YM-254890 on G protein signaling, we tested the impact of YM-254890 on Gs-coupled adenosine A_2_ receptor signaling. As shown in figure 3A, stimulation of HCAEC with 100μM adenosine induced significant elevation of intracellular cAMP as expected. To our surprise, however, treatment of the cells with 30 nM YM-254890 significantly inhibited adenosine A_2_ receptor-induced cAMP production by more than 60% (Fig. 3A), suggesting that YM-254890 may block the Gα_s_ protein. To support this notion, we further studied the impact of YM-254890 on other Gs-coupled receptor-induced cAMP elevation. Figure 3B shows that stimulation of the Gs-coupled β_2_-adrenergic receptors by isoproterenol or the Gs-coupled PGI_2_ receptors by iloprost, all dose-dependently increased cellular cAMP, both of which were significantly suppressed by YM-254890 pre-treatment, suggesting a general inhibition on Gs proteins by YM-254890.

**Figure 3.**
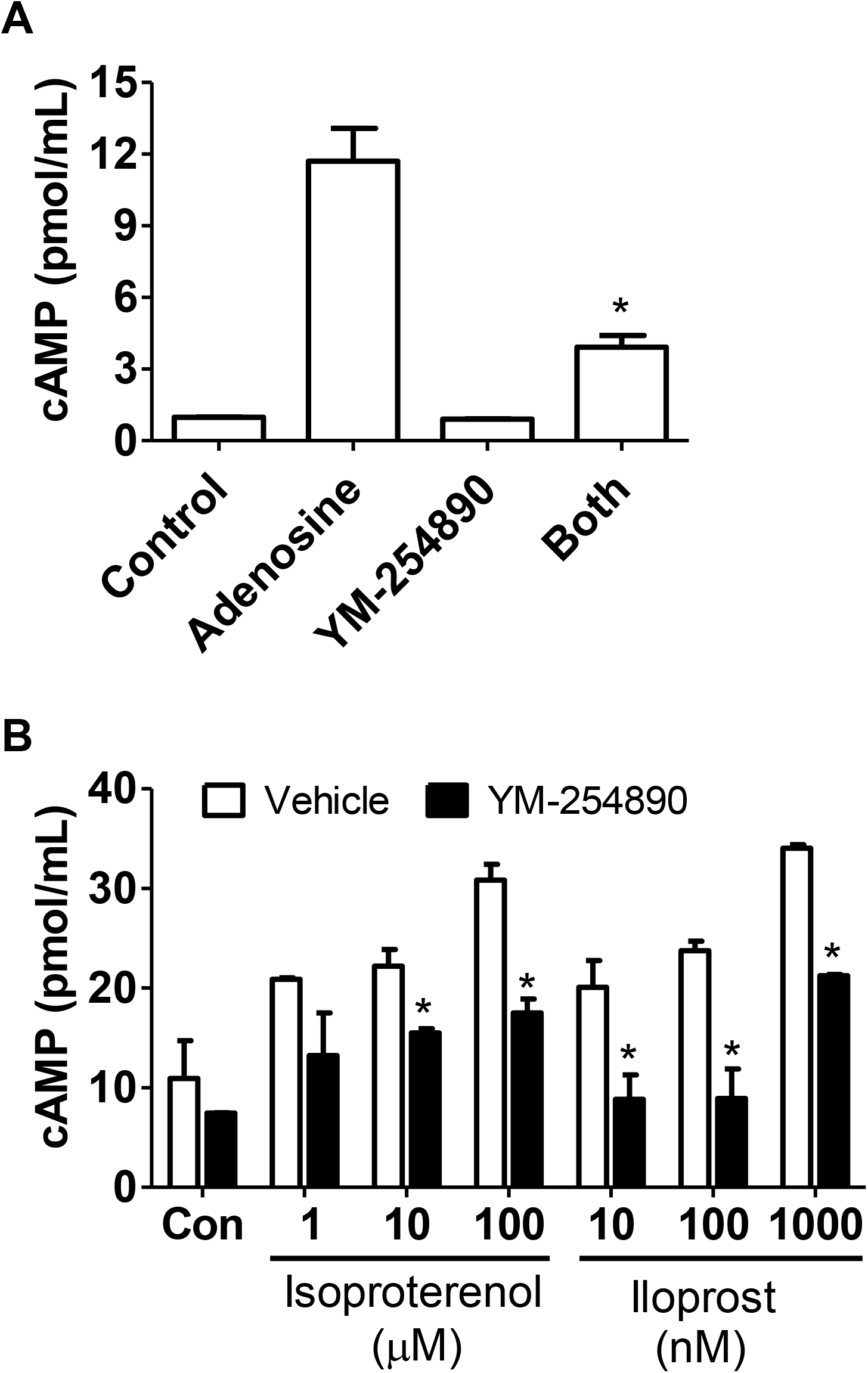
Effect of YM-254890 on Gs-coupled receptor-induced cAMP elevation in HCAEC. HCAEC were pretreated with vehicle or YM-25489 at 30 nM for 40 min before stimulated for 10 min by adenosine (100uM, **A**) or Isoproterenol and Iloptost at different concentrations (**B**), after which cells were lysed for cAMP ELISA assay as detailed in “Materials and methods”. Data are the mean ± SEM from three independent experiments performed in triplicate. * *p*< 0.05 relative to the respective controls.

### Effect of YM-254890 on MAPK signaling triggered by Gq-coupled P2Y2, Gi/o-coupled CXCR4, and multiple Gs-coupled receptors

We then further explored whether YM-254890 would affect MAPK signaling pathways triggered by Gα_q_-coupled P2Y2, Gαi-coupled CXCR4, and Gα_s_-coupled receptors. We pretreated HCAEC with YM-254890 (30 nM) to inhibit the G proteins. UTP and APPNHP (a non-hydrolysable ATP analog) were used to activate the P2Y2R, SDF-1 was used to activate CXCR4-Gi/o protein signaling, and isoproterenol, iloprost or adenosine to activate their respective Gα_s_-coupled receptors. As shown in figure 4, stimulation of the P2Y2R by APPNHP or UTP activated ERK1/2 as evidenced by their increased phosphorylation. YM-254890 treatment abolished P2Y2R-mediated ERK1/2 activation as we expected. Surprisingly, we also found that ERK1/2 activation by isoproterenol-stimulated β_2_ adrenergic receptor or by iloprost-stimulated PGI_2_ receptor were significantly suppressed by YM-254890 as well (Fig. 4B, 4C). Further, YM-254890 pre-treatment also abolished adenosine A2 receptors-mediated ERK1/2 activation (Fig. 4D). However, the biggest surprise to us is that YM-254890 also eliminated SDF-1/CXCR4-mediated ERK1/2 activation in as much as what *PTX* did (Fig. 4E), suggesting that Gi and/or Go proteins may be targeted by YM-254890.

**Figure 4.**
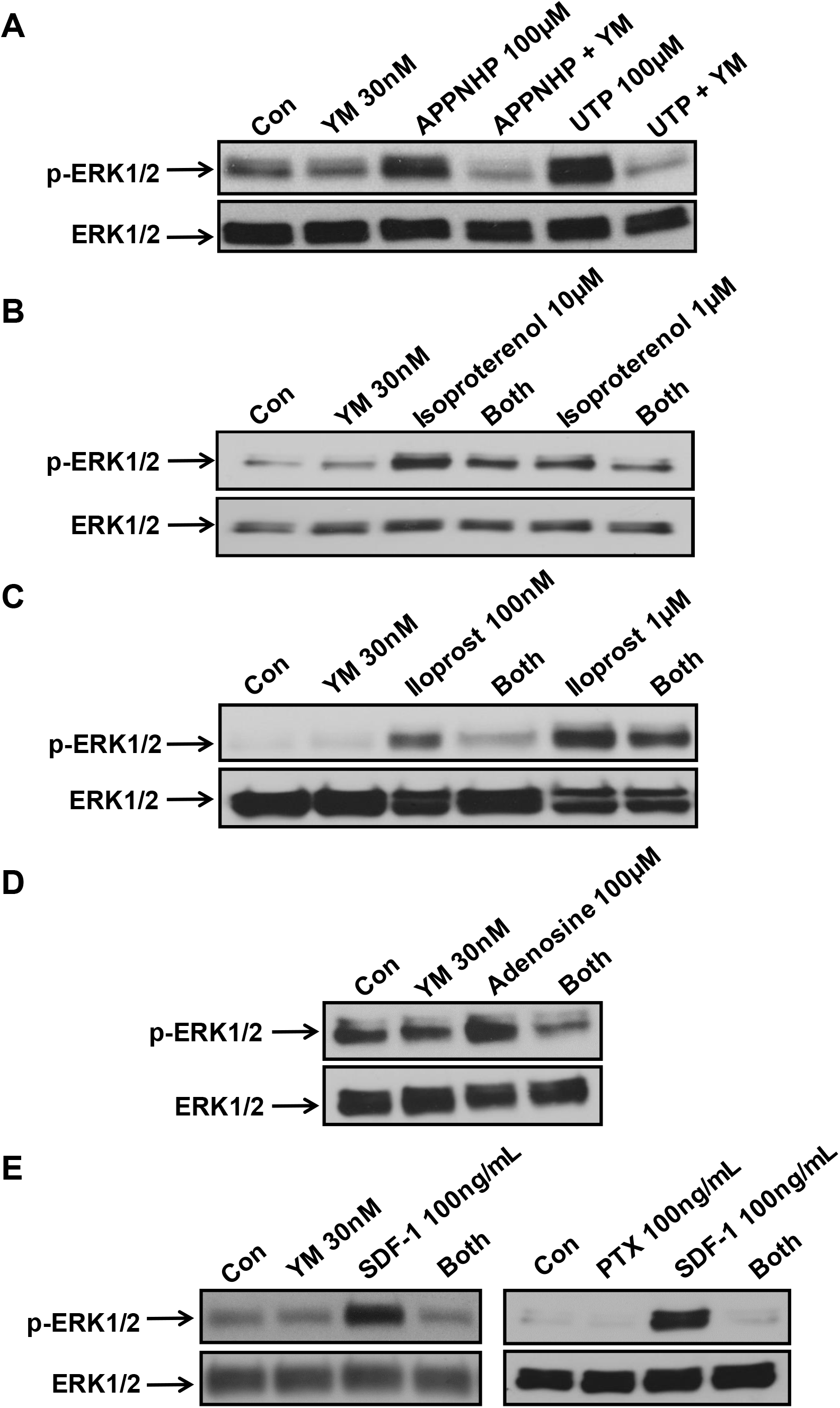
Effect of YM-254890 on MAPK signaling induced by G_q_-coupled P2Y2, G_i_-coupled CXCR4, and several Gs-coupled receptors in HCAEC. Phosphorylation levels of ERK1/2 MAPK were determined by Western blotting assay after HCAEC were stimulated by APPNHP (ATP analog) or UTP for 10 min with or without YM-254890 pretreatment for 40 min (**A**). The same assays were performed for the cells that stimulated with Isoproterenol (**B**), Iloprost (**C**), adenosine (**D**), or SDF-1 (**E**) with or without YM-254890 pretreatment for 40 min. PTX (100 ng/mL) was used as a control to inhibit Gi/o proteins (**E**). Shown are representative data of three independent experiments.

### Effect of YM-254890 on growth factor receptor-mediated cell signaling in HCAEC

Next, we asked whether YM-254890 affects tyrosine kinase receptor-induced cell signaling, which is usually considered G protein-independent. Figure 5 shows that YM-254890 pre-treatment had a negligible to no inhibition on 5% FBS-induced activation of ERK1/2 and JNK (Fig. 5A). Consistently, we also found that YM-254890 did not affect VEGF-induced signaling to the ERK1/2 or JNK pathways (Fig. 5B), indicating that YM-254890 does not impact cell signaling induced by growth factor receptors.

**Figure 5.**
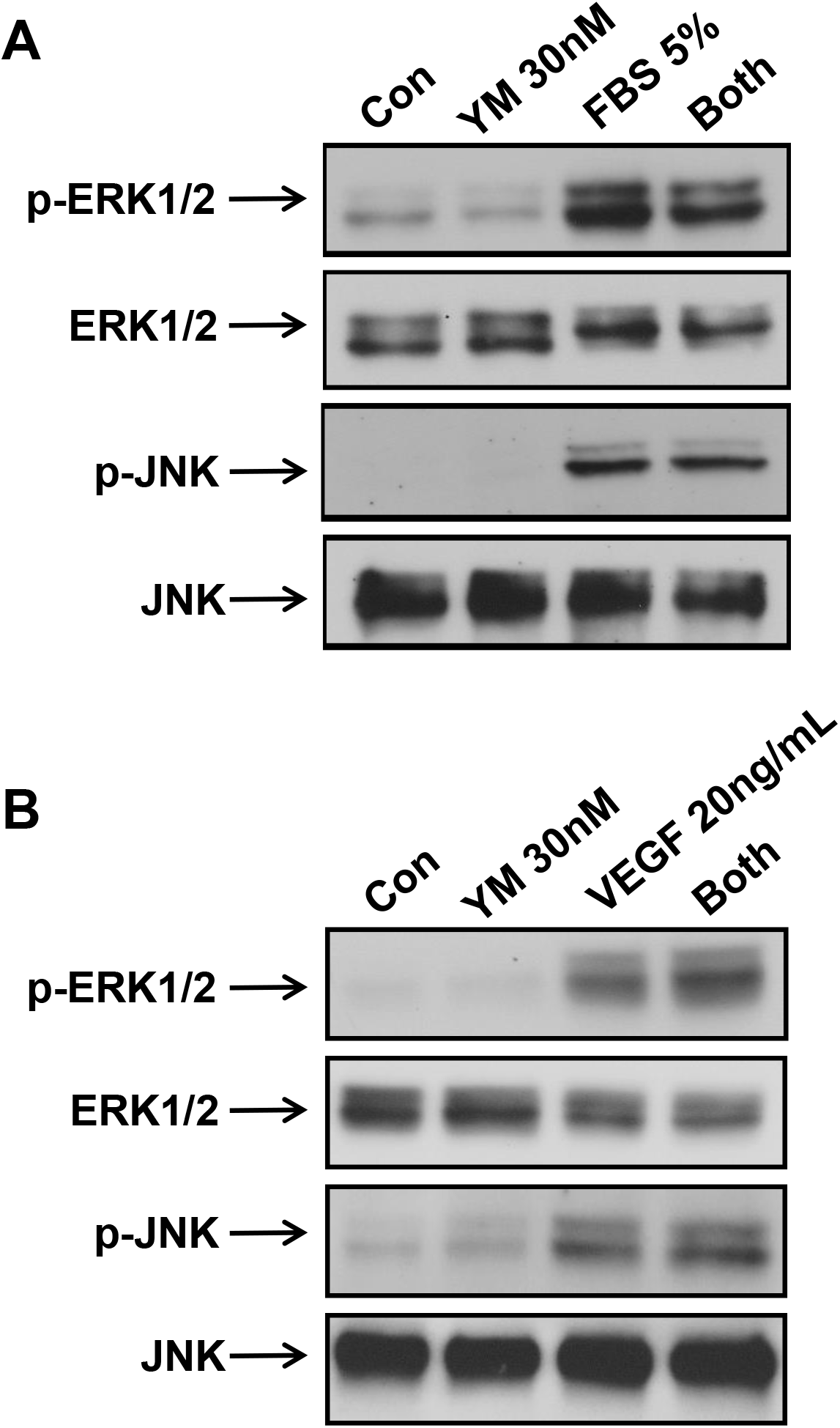
Effect of YM-254890 on growth factor receptor-mediated cell signaling in HCAEC. HCAEC were pretreated with YM-254890 (30 nM) for 40 min before stimulated by 5% FBS (**A**) or 20 ng/mL VEGF (**B**) for 10 min, after which p-ERK1/2 and p-JNK were determined by Western blotting assays. Total ERK and JNK were as loading controls. Shown are representative data of three independent experiments.

### Effect of YM-254890 on non-receptor-mediated cell signaling in HCAEC

To further investigate the specificity of YM-254890, we examined whether YM-254890 would affect non-receptor-mediated cell signaling. As shown in figure 5, stimulation of the HCAEC with the Ca^2+^ ionophore A23187 (17) induced significant Ca^2+^ signaling as expected, which was not significantly affected by YM-254890 pretreatment. Similarly, YM-254890 exhibited no impact on the cellular Ca^2+^ signaling induced by thapsigargin, a selective inhibitor of the sarco/endoplasmic reticulum Ca^2+^ ATPase (Fig. 6B). In addition, stimulation of the cells with forskolin, a direct activator of adenylyl cyclase, significantly increased cellular cAMP level as expected, which was not significantly changed by YM-254890 pre-treatment (Fig. 6C). These data indicate that YM-254890 does not affect non-receptor-mediated Ca^2+^ signaling and cAMP generation in HCAEC.

**Figure 6.**
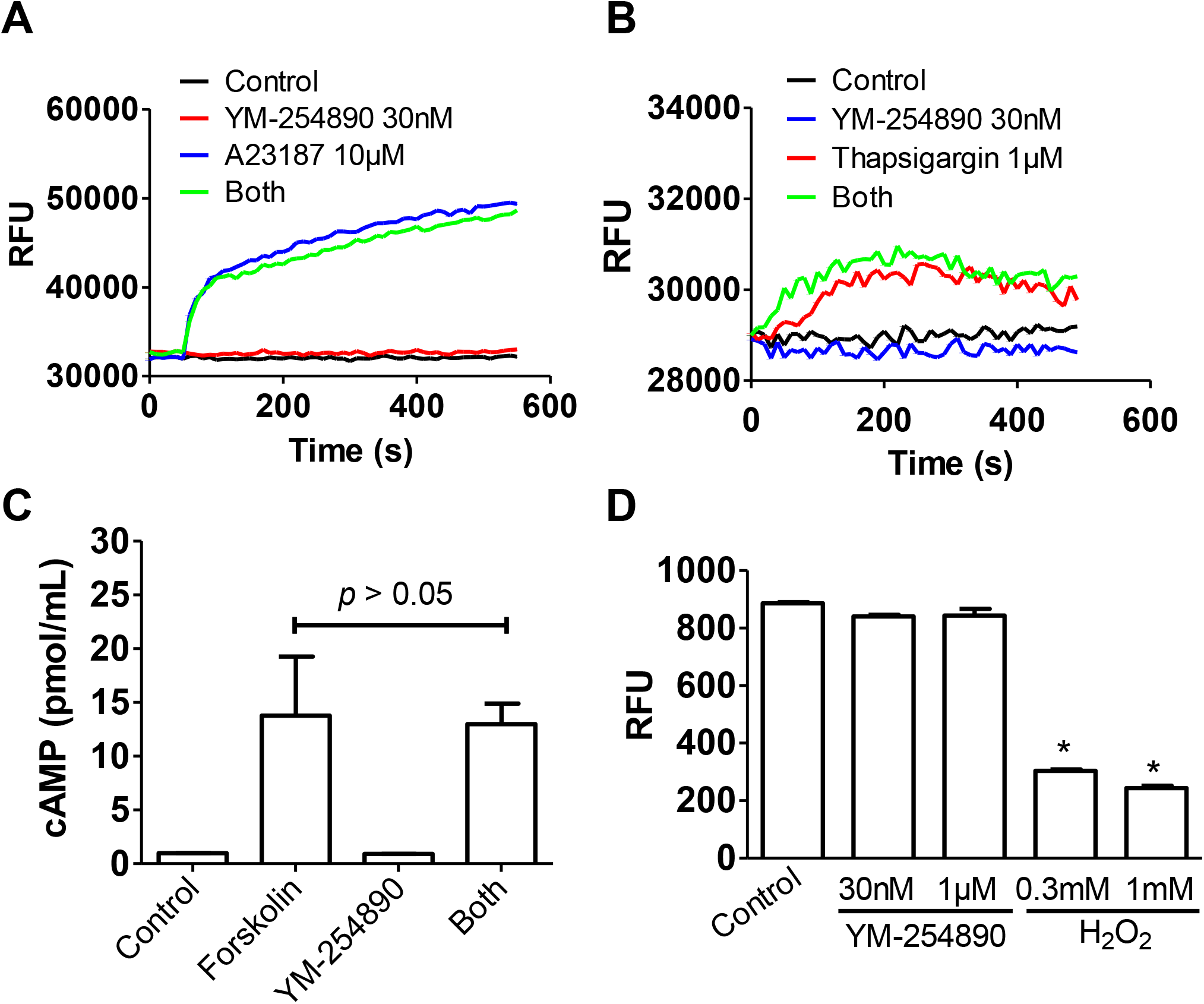
Effect of YM-254890 on non-receptor-mediated cell signaling and its toxicity in HCAEC. A23187 (**A**)- or thapsigargin (**B**)-induced intracellular [Ca^2+^] mobilization were determined in HCAEC after pretreatment of the cells for 40 min with 30 nM YM-254890. Measurement of Ca^2+^ signal was performed by a fluorometer plate reader with a 490/525nm bandpass filter and Fura-4 as the probe, the results of which were shown as relative fluorescence units (RFU). Tracings shown are representative data of three independent experiments. HCAEC were pretreated with vehicle or YM-25489 at 30 nM for 40 min before stimulated for 10 min by forskolin (3 μM), after which cells were lysed for cAMP ELISA assay as detailed in “Materials and methods”. Data are the mean ± SEM from three independent experiments performed in triplicate (**C**). HCAEC were treated with YM-254890 at the indicated concentrations for 60 minutes, after which cellular toxicity was determined by an assay detailed in “Materials and Methods”. H_2_O_2_ was used as a positive control. * *p*< 0.05 relative to the control. n=5.

### No cellular toxicity for YM-254890 in HCAEC

Finally, we tested the overall cellular toxicity of YM-254890. As shown in figure 6D, treatment of the overnight-starved HCAEC with YM-254890 up to 1μM for 1 h showed no significant cellular toxicity, whereas H2O2 treatment at 300 μM or 1 mM all drastically decreased cell viability.

## DISCUSSION

In the present study, we showed for the first time that besides to inhibit Gα_q_ protein signaling, YM-254890 also inhibited Gα_s_ protein signaling in HCAEC. In addition, we are very surprised to find that YM-254890 abolished Gαi protein-mediated MAPK signaling, although it had no effect on Gαi protein-mediated cAMP inhibition, a new pharmacological phenomenon that we termed it “biased inhibition”. Furthermore, we confirmed that YM-254890 had no impact on growth factor receptor-mediated and non-receptor mediated cell signaling in HCAEC. These findings demonstrate that YM-254890 is not a specific Gα_q_ protein inhibitor as previously reported (7, 8); instead, it acts as a mixed inhibitor for multiple G proteins, with a unique feature on Gi/o protein functions.

YM-254890 was originally isolated from the culture broth of Chromobacterium sp. QS3666 as a new anti-platelet agent by Taniguchi M. et al (7, 18, 19). Initial pharmacologic studies indicated that it inhibited ADP-induced platelet aggregation with an IC_50_ below 0.6μM (7). Further mechanistic study showed that it inhibited the Gq-coupled P2Y1 nucleotide receptor-mediated Ca^2+^ mobilization with an IC50 value of 31 nM, without affecting the G_i_-coupled P2Y12 receptor-mediated cAMP suppression up to 40μM (8). In the same study, the research team found that YM-254890 inhibited [^35^S]GTP-γS binding to G_q/11_ proteins in isolated membranes of cells expressing recombinant P2Y1 or muscarinic M1 (both are Gq-coupled), but not the Gi/o-coupled P2Y12 receptors (8), indicating its specificity to the Gq/11 proteins. In the present study, we found that YM-254890 at 30 nM abolished the Ca^2+^ signaling induced by the endogenous P2Y2 receptor, which is the predominant subtype of P2Y nucleotide receptors expressed in human coronary artery EC (13). However, the same concentration of YM-254890 had no impact on A23187- or thapathgagin-induced Ca^2+^ signaling. These results confirmed that YM-254890 acts as a very potent inhibitor for the Gq/11 proteins.

One of the new findings of this study is about the impact of YM-254890 on receptor-mediated Gs protein signaling. As of today, the only available report that showed the functional effect of YM-254890 on Gs-coupled receptor signaling is from the same research team who originally discovered YM-254890 (8). In their original study, Takasaki et al found that YM-254890 up to 10 μM did not affect cAMP accumulation induced by Gs-coupled β_2_-adrenoreceptor activation (8). However, in the present study, we found that YM-254890 at 30 nM blocked adenosine A_2_ receptor-mediated cAMP generation in HCAEC by more than 60%, suggesting potential inhibition of the Gs proteins by YM-254890. To support this notion, we extended our study on other Gs-coupled receptors expressed in HCAEC. Our results clearly showed that YM-254890 at 30 nM, significantly suppressed cAMP accumulation induced by Gs-coupled PGI_2_ receptors and the isoproterenol-activated β_2_-adrenoreceptor as well. Obviously, there is inconsistency between our present finding and the prior study by Takasaki et al regarding the impact of YM-254890 on β_2_-adrenoreceptor cAMP signaling. Although the exact reason for this discrepancy is unknown, we believe that different testing systems may be one of the reasons. In their study, HEK293 cells over-expressed with recombinant β_2_-adrenoreceptor was used for cAMP assay and we determined the effect of YM-254890 on cAMP generation induced by the endogenous β_2_-adrenoreceptor in HCAEC. It is conceivable that a supraphysiological level of β_2_-adrenoreceptors heterologously expressed in HEK293 cells may underestimate the inhibitory effect of YM-254890 on Gs protein-mediated cAMP production. Given that YM-254890 at nano-mole level inhibited multiple Gs-coupled receptors’ signaling to cAMP and the ERK1/2 pathways without affecting forskolin-induced cAMP generation, it is reasonable to conclude that in addition to the Gq proteins, the Gs proteins may be a previously unrecognized target for YM-254890. Although our finding challenged the view of YM-254890 as a selective Gq protein inhibitor, it should be noted that as of today there is no potent Gs protein inhibitor available on the market except for *Cholera toxin* acting as a permanent activator of Gs proteins (20) and few suramin analogs unselectively blocking Gs proteins at >10 μM (21, 22). From this point of view, YM-254890 may be a good parental molecule to be used for further synthesis of highly selective and potent Gs protein blocker for research and maybe clinical use as well. Indeed, a recent elegant in vivo study in mice showed that delivery of YM-254890 caused a marked reduction of blood pressure, accompanying with bradycardia and decreased heart rate (23). These pharmacological actions may not be entirely due to Gq protein blockage by YM-254890 as the authors interpreted; instead, based on the new finding of the present study, it is very likely that the observed depressor action on the heart is due to the previously neglected inhibitory effect of YM-254890 on the Gs proteins which are vital for normal heart functions.

The most interesting and unexpected finding of our present study is that YM-254890 treatment differentially affected multiple signaling pathways mediated by Gi/o proteins. First, we found that treatment of HCAEC with YM-254890 did not affect cAMP reduction induced by Gi/o-coupled CXCR4 receptor. This observation is consistent with all prior reports showing no impact of YM-254890 on the activity of Gi/o proteins activated by other GPCRs (8). However, to our surprise, when we monitored other signaling readouts mediated by CXCR4 receptors, we consistently found that YM-254890 at 30 nM abolished CXCR4-mediated activation of the ERK1/2 pathway, without affecting the same pathway triggered by growth factor receptors. These results indicate that unlike the traditional Gi/o inhibitor *pertussis toxin* which uniformly blocked both the cAMP and the MAPK pathways, YM-254890 exhibited an apparent “biased inhibition” towards the Gi/o proteins. Although the precise mechanism of this new pharmacological phenomenon is unclear and beyond the scope of this study, we speculated that several potential molecular modes could explain such an unusual “biased inhibition”. It has been shown that YM-254890 binds to the Gαq subunit and blocks the Gq protein activity by preventing the exchange of GDP for GTP, thus locking the Gq protein complex in an inactive GDP-bound form (11). However, it remains unknown whether YM-254890 could induce the same binding with the Gi/o proteins. Since cAMP reduction by CXCR4-Gi/o signaling was not affected by YM-254890, we deduced that YM-254890 may not bind to the Gαi subunit. However, it is plausible that YM-254890 may bind and inhibit the Go protein and that the CXCR4 receptor may rely on Go protein to activate the ERK1/2 signaling and the Gi protein is responsible for CXCR4-mediated cAMP reduction, a phenomena that cannot be differentiated by the traditional Gi/o dual inhibitor *pertussis toxin*. Alternatively, there may be a link between full inhibition of Gq protein and “biased inhibition” of Gi/o protein by YM-254890. It was known that YM-254890 stabilizes the Gαq-GDP state, which has a high affinity for Gβγ. Thus, it is conceivable that the YM-254890/Gαq-GDP complex may sequester or “hijack” Gβγ released from the Gi/o proteins, leading to the inhibition of CXCR4 signaling through Gβγ, but not the signaling through Gα_i/o_. This interpretation is consistent with the fact that many Gi/o-coupled receptors rely Gβγ to activate MAPK and AKT pathways. Obviously, additional investigation is warranted for further understanding the molecular mechanism of YM-254890 on Gi/o protein signaling.

One of the limitations of our current study is about the potential impact of YM-254890 on G_12/13_ proteins which was not pursued here. This is primarily since there is no single GPCR in HCAEC that is exclusively coupled to G_12/13_ proteins. In addition, it is well established that G_12/13_ proteins activate Rho signaling through RhoGEF; however, the Rho signaling pathway can be activated by Gq protein as well (2). Thus, it is not practical to do functional study on the effect of YM-254890 on G_12/13_ signaling. Therefore, it remains to be determined whether YM-254890 affects G_12/13_ protein signaling.

In summary, we provided the first functional evidence to indicate that YM-254890 is not a Gq-selective inhibitor as previously reported (7); instead, it acts as a potent inhibitor for both the Gq and Gs proteins. In addition, our findings suggest for the first time that YM-254890 may function as a “biased inhibitor” on Gi/o protein signaling as evidenced by no impact on cAMP generation, but suppressing Gi/o-mediated MAPK signaling. Overall, our study implies that it should be more cautious to interpret data when YM-254890 is used to determine which G protein(s) mediate cell signaling of a receptor of interest. Nevertheless, given the fact that YM-254890 does not affect growth factor-induced or non-receptor-mediated cell signaling, it still is a very useful pharmacological tool in functional study, considering its nanomole affinity towards the G proteins with virtually no toxicity up to 10μM in intact cells. Future study focused on YM-254890 interaction with different purified G proteins is needed to further understand its molecular mechanism.

## Acknowledgments

This work was supported in part by NIH funding 1R01HL125279-01A1 (JS). The content is solely the responsibility of the authors and does not necessarily represent the official views of the National Institutes of Health.

## REFERENCES

1. Sriram K, Insel PA. (2018) G Protein-Coupled Receptors as Targets for Approved Drugs: How Many Targets and How Many Drugs? Mol Pharmacol. 93(4):251–258

2. Campbell AP, Smrcka AV. (2018) Targeting G protein-coupled receptor signalling by blocking G proteins. Nat Rev Drug Discov. 17(11):789–803

3. Neves SR, Ram PT, Iyengar R. (2002) G protein pathways. Science. 296(5573):1636–1639

4. Schlyer S, Horuk R. (2006) I want a new drug: G-protein-coupled receptors in drug development. Drug Discov Today. 11(11-12):481–493

5. Vassilatis DK, Hohmann JG, Zeng H, Li F, Ranchalis JE, Mortrud MT, Brown A, Rodriguez SS, Weller JR, Wright AC, Bergmann JE, Gaitanaris GA. (2003) The G protein-coupled receptor repertoires of human and mouse. Proc Natl Acad Sci U S A. 100(8):4903–4908

6. Locht C, Coutte L, Mielcarek N. (2011) The ins and outs of pertussis toxin. FEBS J. 278(23):4668–4682

7. Taniguchi M, Nagai K, Arao N, Kawasaki T, Saito T, Moritani Y, Takasaki J, Hayashi K, Fujita S, Suzuki K, Tsukamoto S. (2003) YM-254890, a novel platelet aggregation inhibitor produced by Chromobacterium sp. QS3666. J Antibiot (Tokyo). 56(4):358–363

8. Takasaki J, Saito T, Taniguchi M, Kawasaki T, Moritani Y, Hayashi K, Kobori M. (2004) A novel Galphaq/11-selective inhibitor. J Biol Chem. 279(46):47438–47445

9. Gao ZG, Jacobson KA. (2016) On the selectivity of the Gα_q_ inhibitor UBO-QIC: A comparison with the Gαi inhibitor pertussis toxin. Biochem Pharmacol.107:59–66

10. Kukkonen JP. (2016) G-protein inhibition profile of the reported Gq/11 inhibitor UBO-QIC. Biochem Biophys Res Commun. 469(1):101–107

11. Nishimura A, Kitano K, Takasaki J, Taniguchi M, Mizuno N, Tago K, Hakoshima T, Itoh H. (2010) Structural basis for the specific inhibition of heterotrimeric Gq protein by a small molecule. Proc Natl Acad Sci U S A. 107(31):13666–13671

12. Shukla AK, Xiao K, Lefkowitz RJ. (2011) Emerging paradigms of β-arrestin-dependent seven transmembrane receptor signaling. Trends Biochem Sci. 36(9):457–469

13. Ding L, Ma W, Littmann T, Camp R, Shen J. (2011) The P2Y2 nucleotide receptor mediates tissue factor expression in human coronary artery endothelial cells. J Biol Chem. 286(30):27027–27038

14. Liu Y, Zhang L, Wang C, Roy S, Shen J. (2016) Purinergic P2Y2 Receptor Control of Tissue Factor Transcription in Human Coronary Artery Endothelial Cells: NEW AP-1 TRANSCRIPTION FACTOR SITE AND NEGATIVE REGULATOR. J Biol Chem. 291(4):1553–1563

15. Ma W, Liu Y, Wang C, Zhang L, Crocker L, Shen J. (2014) Atorvastatin inhibits CXCR7 induction to reduce macrophage migration. Biochem Pharmacol. 89(1):99–108

16. Shen J, Halenda SP, Sturek M, Wilden PA. (2005) Novel mitogenic effect of adenosine on coronary artery smooth muscle cells: role for the A1 adenosine receptor. Circ Res. 96(9):982–990

17. Abbott BJ, Fukuda DS, Dorman DE, Occolowitz JL, Debono M, Farhner L. (1979) Microbial transformation of A23187, a divalent cation ionophore antibiotic. Antimicrob Agents Chemother. 16(6):808–812

18. Kawasaki T, Taniguchi M, Moritani Y, Uemura T, Shigenaga T, Takamatsu H, Hayashi K, Takasaki J, Saito T, Nagai K. (2005) Pharmacological properties of YM-254890, a specific G(alpha)q/11 inhibitor, on thrombosis and neointima formation in mice. Thromb Haemost. 94(1):184–192

19. Uemura T, Kawasaki T, Taniguchi M, Moritani Y, Hayashi K, Saito T, Takasaki J, Uchida W, Miyata K. (2006) Biological properties of a specific Galpha q/11 inhibitor, YM-254890, on platelet functions and thrombus formation under high-shear stress. Br J Pharmacol. 148(1):61–69

20. Ayoub MA. (2018) Small molecules targeting heterotrimeric G proteins. Eur J Pharmacol.826:169–178

21. Freissmuth M, Boehm S, Beindl W, Nickel P, Ijzerman AP, Hohenegger M, Nanoff C. (1996) Suramin analogues as subtype-selective G protein inhibitors. Mol Pharmacol. 49(4):602–611

22. Hohenegger M, Waldhoer M, Beindl W, Böing B, Kreimeyer A, Nickel P, Nanoff C, Freissmuth M. (1998) Gsalpha-selective G protein antagonists. Proc Natl Acad Sci U S A. 95(1):346–351

23. Meleka MM, Edwards AJ, Xia J, Dahlen SA, Mohanty I, Medcalf M, Aggarwal S, Moeller KD, Mortensen OV, Osei-Owusu P. (2019) Anti-hypertensive mechanisms of cyclic depsipeptide inhibitor ligands for Gq/11 class G proteins. Pharmacol Res.141:264–275

